# How salience enhances inhibitory control: An analysis of electro-cortical mechanisms

**DOI:** 10.1101/2022.01.12.475991

**Authors:** J. Leon Kenemans, Iris Schutte, Sam Van Bijnen, H. N. Alexander Logemann

## Abstract

Stop-signal tasks (SSTs) combined with human electro-cortical recordings (Event-Related Potentials, ERPs) have revealed mechanisms associated with successful stopping (relative to failed), presumably contributing to inhibitory control. The corresponding ERP signatures have been labeled stop N1 (+/- 100-ms latency), stop N2 (200 ms), and stop P3 (160-250 ms), and argued to reflect more proactive (N1) versus more reactive (N2, P3) mechanisms. However, stop N1 and stop N2, as well as latencies of stop-P3, appear to be quite inconsistent across studies. The present work addressed the possible influence of stop-signal salience, expecting high salience to induce clear stop N1s but reduced stop N2s, and short-latency stop P3s.

Three SST varieties were combined with high-resolution EEG. An imperative visual (go) stimulus was occasionally followed by a subsequent (stop) stimulus that signalled to withhold the just initiated response. Stop-Signal Reaction Times (SSRTs) decreased linearly from visual-low to visual-high-salience to auditory. Auditory Stop N1 was replicated. A C1-like visual evoked potential (latency < 100 ms) was observed only with high salience, but not robustly associated with successful versus failed stops. Using the successful-failed contrast a visual stop-N1 analogue (112-156 ms post-stop-signal) was identified, as was right-frontal stop N2, but neither was sensitive to salience. Stop P3 had shorter latency for high than for low salience, and the extent of the early high-salience stop P3 correlated inversely with SSRT. These results suggest that salience-enhanced inhibitory control as manifest in SSRTs is associated with reactive rather than proactive electrocortical mechanisms.

## Introduction

Inhibitory control is crucial in everyday functioning and deficits in inhibitory control are implicated in disorders such as Attention Deficit / Hyperactivity Disorder (e.g., Kenemans et al., 2005). Inhibitory motor control has been extensively studied using the Stop-Signal Task (SST) in combination with electroencephalography (EEG) (De Jong, Coles, Logan, & Gratton, 1990). In the classical auditory-stop SST variety, participants are presented with a visual choice (go) stimulus to which they have to respond by means of pressing one of two buttons. For a minority of trials the go stimulus is followed by an auditory stop stimulus, and participants are required to suppress their ongoing response. This ‘stopping’ ability is preferably qyantified as Stop Signal Reaction Time (SSRT), a generally accepted behavioral measure of inhibitory motor control (De Jong, Coles, & Logan, 1995; De Jong et al., 1990; Verbruggen et al., 2019).

Previous studies have shown that successful, relative to failed, stopping is associated with an enhanced response in the auditory cortex to the auditory stop signal (Bekker, Kenemans, Hoeksma, Talsma, & Verbaten, 2005; Bekker, Overtoom, et al., 2005; Hughes et al., 2012; Lansbergen, Bocker, Bekker, & Kenemans, 2007; Overtoom et al., 2009; Matzke et al., 2017; Skippen et al., 2020). This response has the form of a negative brain potential emerging at about 100 ms post-stop-signal (‘stop N1’, the difference in N1 amplitude between successful and failed stops). Another line of evidence has revealed an inverse correlation between extent of damage in right frontal cortex (RFC) and stopping performance (Aron, Fletcher, Bullmore, Sahakian, & Robbins, 2003). These observations are consistent with a model in which a potentiated inhibitory link between motor and auditory cortex is under the control of (a tonically active) RFC (Overtoom et al., 2009). Furthermore, ADHD patients, characterized by excessive impulsivity, lack the stop N1 effect in the auditory SST, while their standard medication, methylphenidate, restores their stop N1 (Bekker et al., 2005; Overtoom et al., 2009). An extensive review of this presumably proactive inhibitory mechanism, as well as of its reactive counterpart, is provided by Kenemans (2015).

For stop signals presented in the same visual modality as the go stimuli the situation is less clear. To our knowledge, a visual analogue (posterior scalp distribution, latency < 200ms) of the stop N1 has been reported only in one MEG-SST study (Boehler et al., 2009). In other studies a visual analogue of the stop N1 has not been reported; instead, researchers observed a longer-latency electrocortical component which is generally absent following auditory stop signals. This ‘stop N2’ is a right-frontal negative wave at about 200-ms latency, and is also associated with successful (relative to failed) stopping (Schmajuk, Liotti, Busse, & Woldorff, 2006). Furthermore, ADHD patients present with a smaller N2 effect (successful versus failed, or stop N2) in the visual SST which is reversed by methylphenidate (Pliszka et al., 2007; Pliszka, Liotti, & Woldorff, 2000).

Assuming that the stop N2 indeed reflects right-hemisphere frontal activation, we argue that the RFC contributes to stopping performance in different SST varieties but in different manners (Kenemans, 2015). In the auditory SST and some visual SSTs, it proactively initiates and maintains the potentiation of an inhibitory connection between auditory cortex and the motor system. In other visual SSTs, the RFC is recruited reactively only after the presentation of the stop signal, and then contributes to successful stopping by directly signaling to the motor system. This model is consistent with the effect of RFC damage on stopping performance in the auditory SST (where no stop N2 is evident), as well as with deficiencies of stopping with ADHD, and remediation by methylphenidate, in both auditory and visual SSTs (assuming that in both cases RFC functioning is deficient but augmented by methylphenidate).

Here we address the question whether stop N1 and stop N2 can really be dissociated when scrutinized in one sample of subjects, using SST varieties expected to activate either stop N1 or stop N2. In addition, we consider the role of stop-signal salience. Especially auditory stop signals, against a background of visual go stimuli, can be viewed as having a high level of salience, as compared to visual stop signals with properties very similar to the visual go stimuli (e.g., both classes consist of letter symbols, as in Schmajuk et al., 2006). The fact that such low-salient stop signals have been associated mainly with stop N2, and auditory stop signals with stop N1, is consistent with the hypothesis that proactive recruitment of RFC depends on salience. This may sound contra-intuitive, but please note that proactive anticipation of a stop signal may occur more readilty when the stop signal stands out more clearly from the context of the go stimuli.

If salience is crucial for dissociating stop N1 and stop N2, then this should also be manifest within the visual modality. In the present study, we used letter symbols as go stimuli; for stop signals either a low-salient (a dollar sign) or a high-salient stimulus (red flag) was used (Figure 1b and c). For high salience, a visual analogue of the auditory (Figure 1a) stop N1 was expected, and for low salience a stop N2. In addition, we expected SSRTs to decrease from visual-low to visual-high salience to auditory; this is because if salience facilitates stopping mechanisms, this should translate into a similar effect on stopping performance (Blizzard et al., 2016; Montanari et al., 2017; Van der Schoot et al., 2005). As a salience-manipulation check, we evaluated short-latency (<100 ms) visual evoked potentials (VEPs) for the higher impact of high-salient vs. low-salient stop signals in visual cortex.

**Figure 1.**
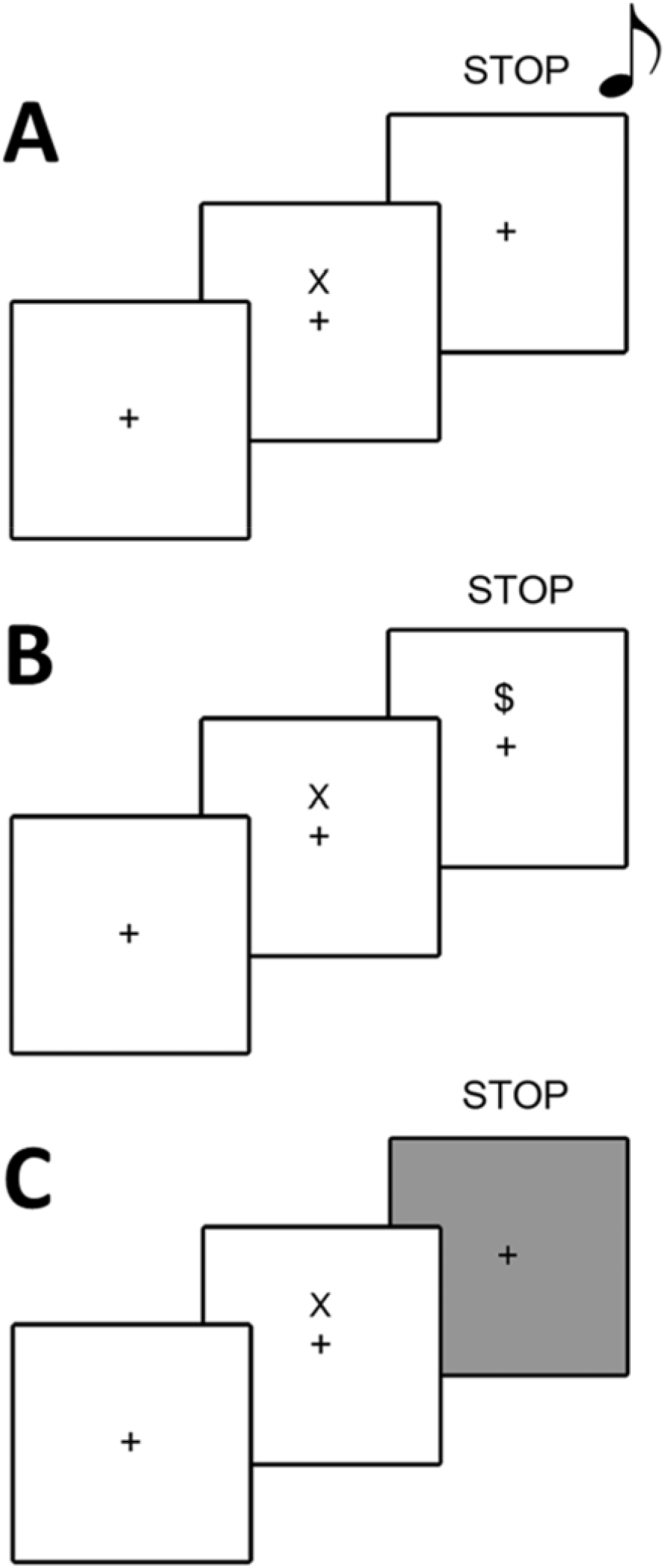
Three SST varieties: A) Auditory stop signal; B) Low-salient visual stop signal; C) High-salient visual stop signal (red screen).

Yet another cortical correlate of successful stopping, is the ‘stop P3’ (De Jong et al., 1990). There is strong evidence that the stop P3 is actually a more generic frontal P3 (fP3) that reflects a general reactive behavioral-interrupt mechanism, implemented in dorsal-medial prefrontal cortex, and activated by salient or otherwise potentially relevant stimuli (Kenemans, 2015; Polich, 2007; Wessel & Aron, 2013, 2017). The existence of this reactive correlate of successful stopping is theoretically important. For example, with auditory stop signals, in ADHD, stopping is slower but nevertheless successful, while there is no stop-N1 effect. The stop-P3 effect however is significant in ADHD, so a cortical correlate of successful stopping can still be demonstrated. Because SSRTs are generally longer with visual than with auditory stop signals, and stop P3 latency is correlated with SSRT (Wessel & Aron, 2014), we expected longer stop-P3 latencies for visual than for auditory stop signals. In addition, it was expected that both SSRT and stop-P3 latencies would be shorter for high, relative to low visual salience.

## Method

### Participants

Based on Bekker et al. (2004-stop N1), Schmajuk et al. (2006-stop N2), and Boehler et al. (2008-visual stop N1), N=20 should be sufficient to detect stop N1, stop N2, and visual-stop N1 effects. However, it was anticipated that to detect the crucial interaction effect (visual stop-N1 present in high-salience setup (Figure 1c) but not in low salience setup (Figure 1b)), a larger sample might be necessary. Therefore, in the ethical-approval protocol it was specified (before the experiment had started) that after having acquired full data sets for 20 participants, an interim analysis of the crucial interaction effect would be conducted. If the interim interaction effect would not be significant at p <.05 whereas there was a clear indication of a sizable stop N1 for high-salient but not for low-salient, then additional participants would be recruited, up to a maximum of 50 in total. After having collected data for 21 initial participants, plus 10 additional participants, due to practical circumstances (personnel dropout) data collection was halted. For the visual-stop conditions, for two participants EEG data were lost or not usable, yielding a final total N of 29. Inspection of the Bayes factors for the likelihood of the data given the null hypothesis, relative to that given the alternative hypothesis (BF01 for the crucial interaction, JASP v0.14, Goss-Sampson et al., 2020), revealed only an increase in BF01 from N=20 to N=29, rendering it unlikely that adding data of further participants would increase the relative likelihood for the alternative hypothesis (see Results for details). Therefore, the eventual sample consisted of 31 healthy participants (19 female) recruited from the student population at Utrecht University, The Netherlands. Mean age was 22.9 years (SD = 3.3, range 19-31 years; age data for 3 subjects lost). All subjects claimed to have normal hearing and normal or corrected-to-normal vision and signed an informed consent written in accordance with the guidelines of, and approved by, the Ethics Committee of the University Medical Centre Utrecht. Participants were paid euro 21,-for their participation.

### Procedure

After participants signed the informed consent, they provided standard demographic information and the EEG cap was placed. They were seated in a dark sound-attenuated room, approximately 90cm from the computer screen. Three conditions of the SST, which were counterbalanced across participants, were performed. Participants were instructed to respond as quickly and accurately as possible to the Go stimuli but to refrain from responding after presentation of a stop signal. After each experimental block a comparison was made between the mean RT to Go stimuli and to the Go stimuli of the practice block. If participants slowed down more than 1.5 times, they were instructed to speed up, and they were instructed to slow down if less than 40 percent successful stops were made. Halfway through the experiment participants were offered a break of about 15 minutes.

#### The Stop Signal Task (SST)

A generic SST was modeled after a previously reported study reporting on the N2 effect (Schmajuk et al., 2006). The primary task consisted of a two-choice response task. In this task participants had to discriminate between two go-stimuli, the letters “X” and “O” (visual angle for both letters, 1.6° x 1.6°), by means of pressing either the left or the right button with respectively the left or the right index finger. Go stimuli were 150ms in duration and were presented sequentially slightly above a central persistent fixation cross. The trial-to-trial interval was varied between 1.5 to 1.8 seconds to prevent expectation effects.

On 25 percent of the trials, a stop signal was presented after the go stimulus (Figure 1). The stop signal characteristic was varied across three conditions. In the auditory condition, the stop signal consisted of a 1000hz-72dB tone, presented binaurally for 150ms through in-ear headphones. In the visual high-salience condition the stop signal was a red background of the computer screen, presented for 150 ms. The stop signal in the visual low-salience condition was a ‘$’ sign (equal visual angle as go-stimuli), presented for 150 ms at the same location as the go stimuli. The experiment consisted of a total of 25 blocks. The first block consisted of 126 go trials, and was used as a practice block and to establish a baseline reaction time (RT). Subsequently, the three stop signal conditions were presented (as said, order counterbalanced across participants).

Each condition started with a base block, used to estimate a go-stop interval (stimulus-onset asynchrony or SOA between go-stimulus onset and stop-stimulus onset) that would yield an approximately 50 % stop rate in the subsequent block. The base block was followed by three experimental blocks, each consisting of 128 trials. After these blocks, the finger-stimulus assignment was switched and another base block and three experimental blocks followed. During the very first base block the initial SOA was 250 ms; for the other 5 base blocks the initial SOA was equal to the average SOA of the preceding experimental block, after applying a tracking algorithm (De Jong et al., 1995). This initial SOA was subsequently increased by 50 ms after a successful stop, and shortened by 50 ms after a failed stop trial, while maintaining a minimum of SOA of 250 ms. The resulting final value of each base block was set to be the average SOA for the subsequent experimental block. The average SOA for each second and third experimental block was determined by a tracking algorithm (De Jong et al., 1995), again to yield an approximate stop rate of 50 %. Finally, for each experimental block, SOAs were jittered over 99ms before and after the average SOA for that block. Furthermore, for all base and experimental blocks, the initial or the average SOA was forced to a minimum of 250 ms. Given go-stimulus duration and maximum jitter value, this ensured that there was no temporal overlap between go and stop stimuli.

During calibration after the first 21 participants had been ran, we found a lag between the audio marker and the physical audio signal output. Oscilloscopy revealed that the time difference between marker output from LTP port and the physical audio signal was on average around 50 milliseconds. This was confirmed by the initial estimates of the N1 peak in the auditory ERP, which is normally at 100 ms, but was now at 150 ms (note: the same held for the stop N1 effect). This means that on average the audio stimuli had a 50 millisecond delay. In the analysis of performance and ERPs and performance, a corresponding 50-ms correction was maintained for these participants.

#### EEG data acquisition

EEG signals were recorded with the Active-Two system (Biosemi, Amsterdam, The Netherlands) with 64 Ag-AgCl electrodes. Recording electrodes were placed according to the 10/10 system. EOG electrodes were placed above and below the left eye and at the outer canthi of both eyes. EEG signals were online referenced to the Common Mode Sense/Driven Right Leg electrode, sampled at 2048 Hz and online low pass filtered at DC to 400 Hz.

### Data analysis

#### Performance analysis

The SSRT was estimated as follows. The proportion of successful stops (stop trials with no responses) was calculated and corrected for omissions using the Tannock correction (Tannock, Schachar, Carr, Chajczyk, & Logan, 1989). RTs of valid responses (within 150ms and 1500ms post stimulus onset) on go trials were rank-ordered from shortest to longest. The N^th^ RT to go stimuli was calculated by multiplying the total number of RTs with 1-(corrected proportion of inhibition). SSRT was estimated by taking the N^th^ RT minus the average go-stop SOA in the block. This is effectively equivalent to the integration method for SSRT estimation (Verbruggen et al., 2019). SSRT, RT, go-stop SOA, and successful-stop rate were statistically analyzed in a standard SPSS/ GLM Repeated Measurs design with one factor Stop-task variety (3 levels).

#### EEG/ERP analysis

EEG was analyzed using Brainvision Analyzer (Brain Products GmbH). Signals were re-referenced to the right mastoid, and down sampled offline to 250 Hz (including automatic anti-aliasing low-pass filtering). Subsequently, a high pass filter of 2 Hz, 24 dB/oct, notch filter of 50 Hz, and low pass filter of 30 Hz, 12 dB/oct were applied. Automatic ocular correction was performed (Gratton, Coles, & Donchin, 1983). After ocular correction, automatic artifact rejection was used (maximal allowed absolute difference between two values: 100 µV, lowest allowed activity within a 100 ms interval: 0.5 µV). Only trials containing a stop-signal were analyzed, separately for successful and for failed stops. Individual ERPs were averaged time-locked to the go and stop stimulus, respectively. These averages were entered into the Adjacent Response Filtering (Adjar) level 2 (Woldorff, 1993) procedure to remove overlap from go ERPs on stop ERPs For the resulting stop ERPs the baseline was set at 0-50ms to remove possible residual overlap. For one subject in all conditions, for a second in specifically the auditory, and for a third one in specifically the visual conditions, EEG data were lost due to technical problems or excessive artifacts. Hence for all conditions 29 participants remained in the EEG analyses (in contrast to 31 for performance analysis).

For the auditory condition the Adjar procedure yielded huge artifacts in the baseline for two participants, although N1 amplitudes were very much comparable before and after Adjar application, and the non-Adjared data were eventually used specifically for these two participant in the auditory condition.

#### Auditory stop N1, VEP, stop P3

Analysis of auditory stop N1 proceeded along the lines of our earlier work (Bekker et al., 2005ab; Lansbergen et al., 2007; Overtoom et al., 2009). As to the VEP, preluding on the results, a pronounced C1-like deflection between 50 and 100 ms was quantified as the average amplitude in the 64-80-ms-latency interval. For stop P3, onset latencies were expected between 160 and 200 ms (auditory), 220-250 ms (low visual salience; Wessel & Aron, 2014), and around 200 ms (high visual salience); analysis proceeded in terms of both onset and peak latency. P3-peak latencies were quantified in the 132-360-ms latency window, separately for the auditory and the two visual conditions.

With respect to the VEP and visual-stop P3, unless stated otherwise, the statistical design for all average-amplitude measures in specific time windows consisted of a Salience (2) x Stop-success (2) setup in the context of a standard SPSS GLM/ Repeated Measures, or using custom made software. In addition, Bayes factors were calculated in case of non-significant Salience x Stop-success interaction or Stop-success main effects.

Recently significant correlations between N1 and P3 amplitude, separately for successful and failed stops, on the one hand, and SSRT on the other have been reported (Skippen et al., 2020), prompting us to attempt replication of these results, which would provide support for a functional relationship between N1/ P3 and SSRT. As our general approach to these functional relations concerns stop N1 and stop P3 rather than N1 and P3 for successful and failed separately, we explored these correlations with SSRT also.

#### Search for visual stop-N1 analogue

As mentioned there is only one core reference to guide the search for the visual stop N1 (Boehler et al., 2009). These authors reported a stronger event-related field (ERF) between 100-200 ms latency for successful compared to failed stops. Their intracranial-source approximation included generators in posterior cingulate gyrus (PCG) and in bilateral occipital-temporal cortex. Based on this, we designed a dual search space consisting of (1) an array of posterior midline electrodes (Iz up to Cz) and (2) bilateral PO7 and PO8 locations. Given the relatively deep medial PCG source, we anticipated across-midline-electrode patterns of a constant, linear and/ or quadratic form. This was translated into a statistical design which tested stop-success main effects and salience-dependent stop-success effects in interaction with polynomial 0-order (average), linear and quadratic trend effects. This design was tested separately for adjacent 20-ms windows ranging from 68 to 212 ms latency post-visual-stop-signal, so as to include both the C1 and the stop N2 window.

These midline-electrode design settings were supplemented with a similar design in which the polynomial terms were replaced with PO7 and PO8 electrode values as a third factor. In the complete midline-lateral design, to account for Type-I error inflation, a critical alpha of .001 was maintained.

Also here, Bayes factors were calculated in case of non-significant Salience x Stop-success interaction or Stop-success main effects.

#### Search for visual stop-N2

To test for the presence of the visual stop-N2, we adopted the latency window (196-212 ms) and the region of interest (electrodes F6, FC6, and F8) as specified by Schmajuk et al. (2006). Average amplitudes across this spatial-temporal window were tested for a main effect of stopping success, and for the interaction between the effects of stopping success and salience. Also here, a critical alpha of .001 was maintained. Again, Bayes factors were calculated in case of non-significant Salience x Stop-success interaction or Stop-success main effects.

#### Further additional baseline correction

A control analysis for the Salience x Stop success design revealed, for the visual conditions, Stop-success effects in the interval from -100 to 0 ms relative to stop-signal onset (pre-stimulus interval). This suggests that in spite of the Adjar procedure, activity induced by the Go stimuli that differed as a function of Stop success was still contaminating the estimation of the stop-signal ERPs. This prompted us to apply baseline criteria that were even more strict than the initially defined average values in the 0-50 ms post-stop-signal interval. These additional baseline criteria will be specified in the Results section.

## Results

### Performance

Stopping speed as indexed by SSRT increased from the auditory condition to the visual high-salience condition to the visual low-salience condition (181.7 ms to 202.6 ms to 232.7 ms; F(2,29) = 28.6, *p* < 0.001). SSRT was significantly longer for low as opposed to high visual salience (F(1,30) = 24.3, *p* < 0.001), and was significantly longer for the latter compared to auditory (F(1,30) = 15.7, *p* < 0.001). Go RTs were not significantly different between modality conditions (662.3, 667.6, and 687.1 ms, respectively), and neither were the SOAs (482.6, 447.0, and 449.3, respectively). Average stop rates ranged from 48% (auditory) to 54.2% (visual high) to 51% (visual low-salient).

### EEG

#### Stop N1

Previously reported results were replicated. Auditory stop signals elicited an N1 that was larger for successful than for failed stops (‘stop N1’, depicted in Figure 2, with a central distribution). This was statistically confirmed by quantifying the N1 as the average amplitude in the 78-122-ms latency window at Cz, and comparing successful versus failed stop trials (F(1,28) = 35.6, *p <* .*0001*). Inspection of single-subject values revealed that all but three subjects had larger N1s for successful than for failed stops (as visible in Figure 2, left lower inset).

**Figure 2.**
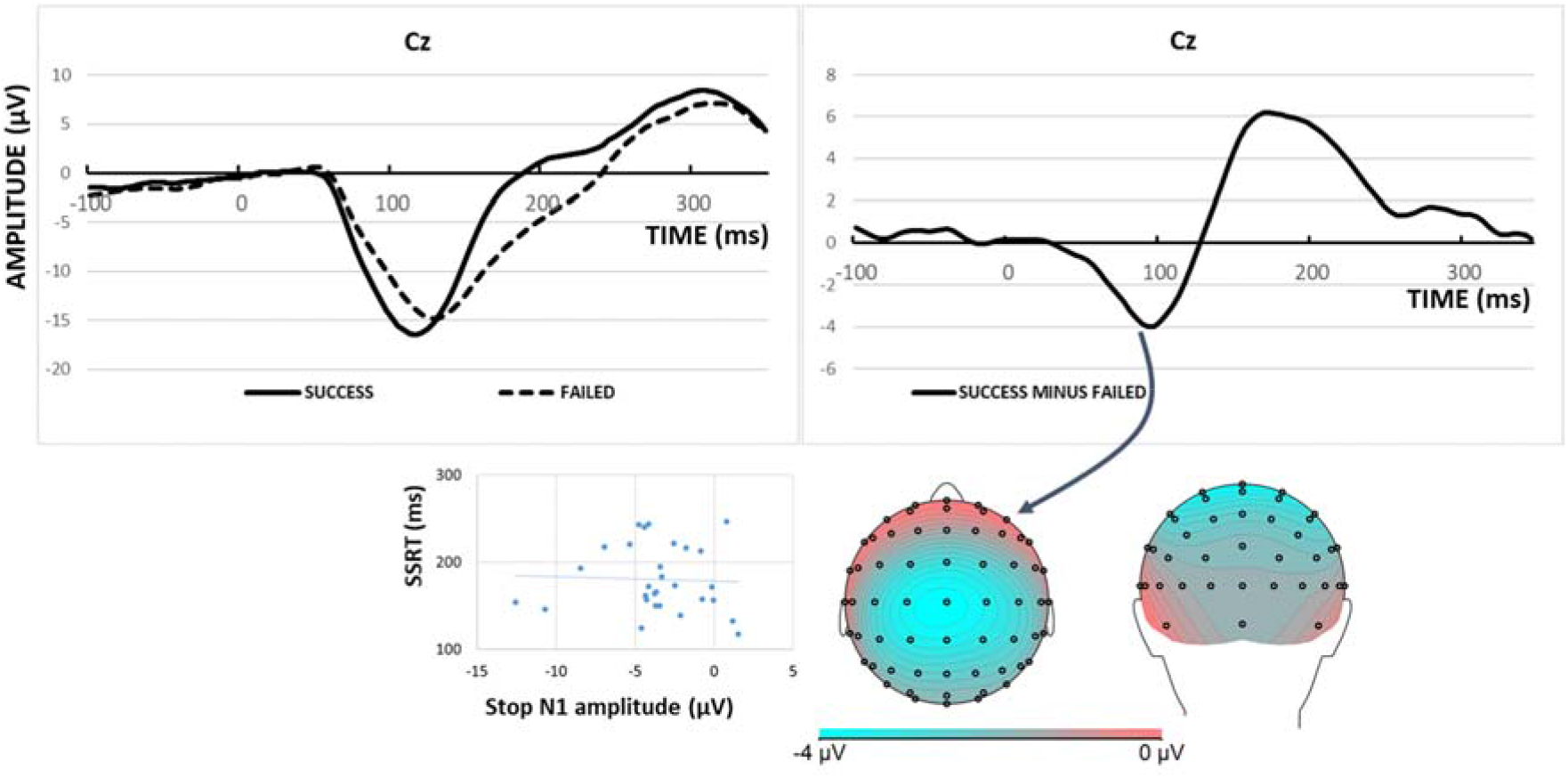
At electrode Cz: the Stop N1 (Successful versus Failed stop), and scalp distribution (92-112 ms) for the auditory condition. Left lower inset: Non-significant correlation between stop N1 (78-122 ms) and SSRT.

Recently significant correlations between N1 amplitude separately for successful and failed stops and SSRT have been reported (Skippen et al., 2020). Attempting to replicate these results for the present data, quite similar, slightly higher, correlation values as in Skippen et al. were observed: Pearson’s r= 0.30 for successful stops and SSRT (*p =* .12; average activity 78-122 post-stimulus) and r= 0.37 for failed stops (*p* < .05). Note that the partial lack of significance at alpha = .05 merely reflects the relatively small present sample size (n=29; n= 156 in Skippen et al.); and that the positive sign of the correlations reflects that bigger negative N1 amplitudes are associated with shorter SSRTs. However, for the stop N1 proper the correlation with SSRT was not significant (Pearson’s r = -.044; see Figure 2, left lower inset).

#### Visual evoked potentials (VEPs)

As can be seen in Figure 3, a C1-like peak was clearly present at medial-occipital sites, between 50 and 100 ms latency, only for salient visual stop signals (main effect of Salience at Oz, 64-80 ms: F(1,28) = 104.1, *p* < .0001). Although the differences were small (see Figure 3), also the effect of Stop success was significant using the 0-50-ms baseline. However, this effect disappeared when the average value in the 50-64-ms window was used as baseline (F(1,28) = 2.2, *p* = .15), whereas the Salience effect was still retained (F(1,28) = 88.2, *p* < .0001). The Bayes factor (BF10) for the alternative hypothesis of an effect of Stop success was 0.53, suggesting anecdotal (inconclusive) evidence in favor of the null hypothesis. With neither baseline there was a significant interaction between Salience and Stop success (BF10 for the 50-64-ms baseline = 0.25, moderate evidence for the null hypothesis).

**Figure 3.**
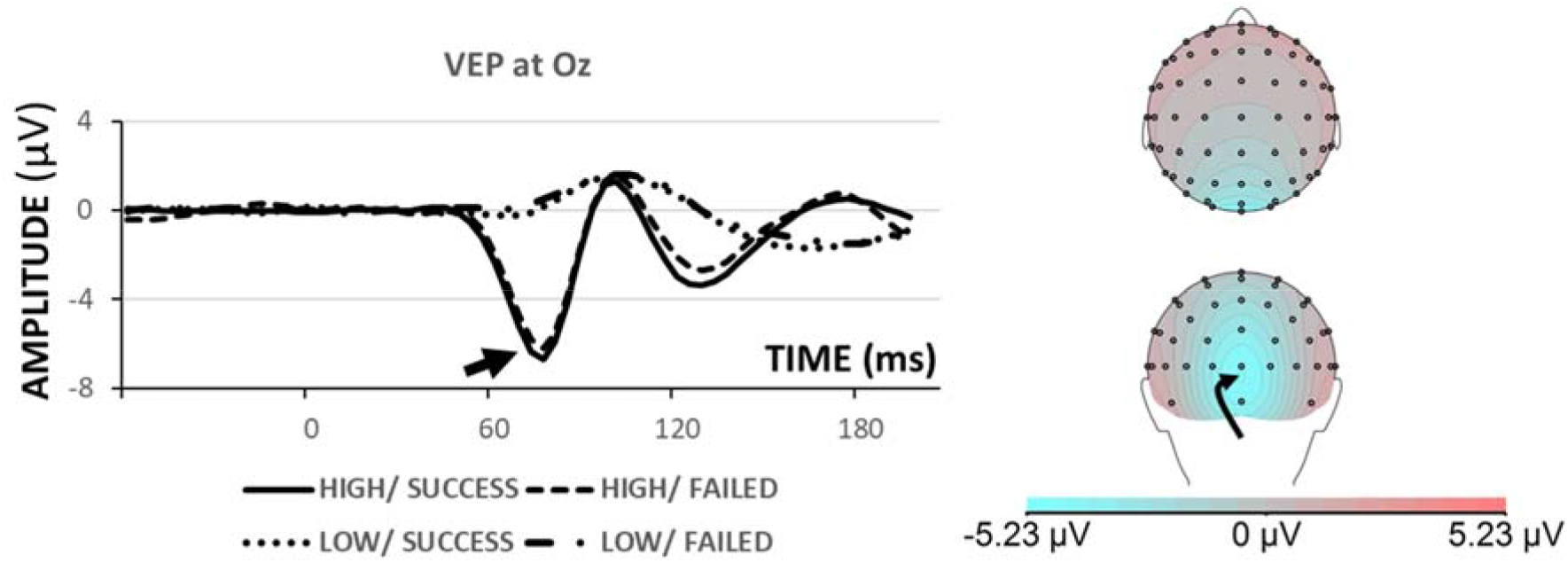
At electrode Oz (right arrow): the C1-like VEP (left arrow), and scalp distribution (64-80 ms) for the four visual conditions.

#### Visual stop-N1 analogue

The “*Search for visual stop-N1 analogue”* procedure outlined in the Method section revealed Stop-Success main effects for the Cz-Iz midline array in the 112-132-ms and 136-152-ms time intervals. These effects were manifest as Stop-Success effects across the midline electrodes on average (0-order), as well as in the form of a linear (1-order) trend across midline electrodes (112-132 ms: F(1,28) = 21.0, *p* = .0001, and 13.5, *p* = .001, respectively; 136-152 ms: F(1,28) = 15.5, *p* = .0005, and 16.2, *p* = .0004, respectively). As can be seen in Figure 4, in both Salience conditions the Successful-versus-Failed contrast revealed a negativity between 100 and 160 ms (that subsequently reversed into a positivity), with a central-parietal scalp distribution. With alpha set at .001, there were no significant interactions between Salience and Stop Success; BF10 for 112-132 ms = 0.25 (0-order), and 0.22 (linear); for 136-152 ms 0.28 and 0.20, respectively; all moderate evidence for the null hypothesis. At alpha = .001 there was also no effect involving Stop Success at the PO7/ PO8 sites.

**Figure 4.**
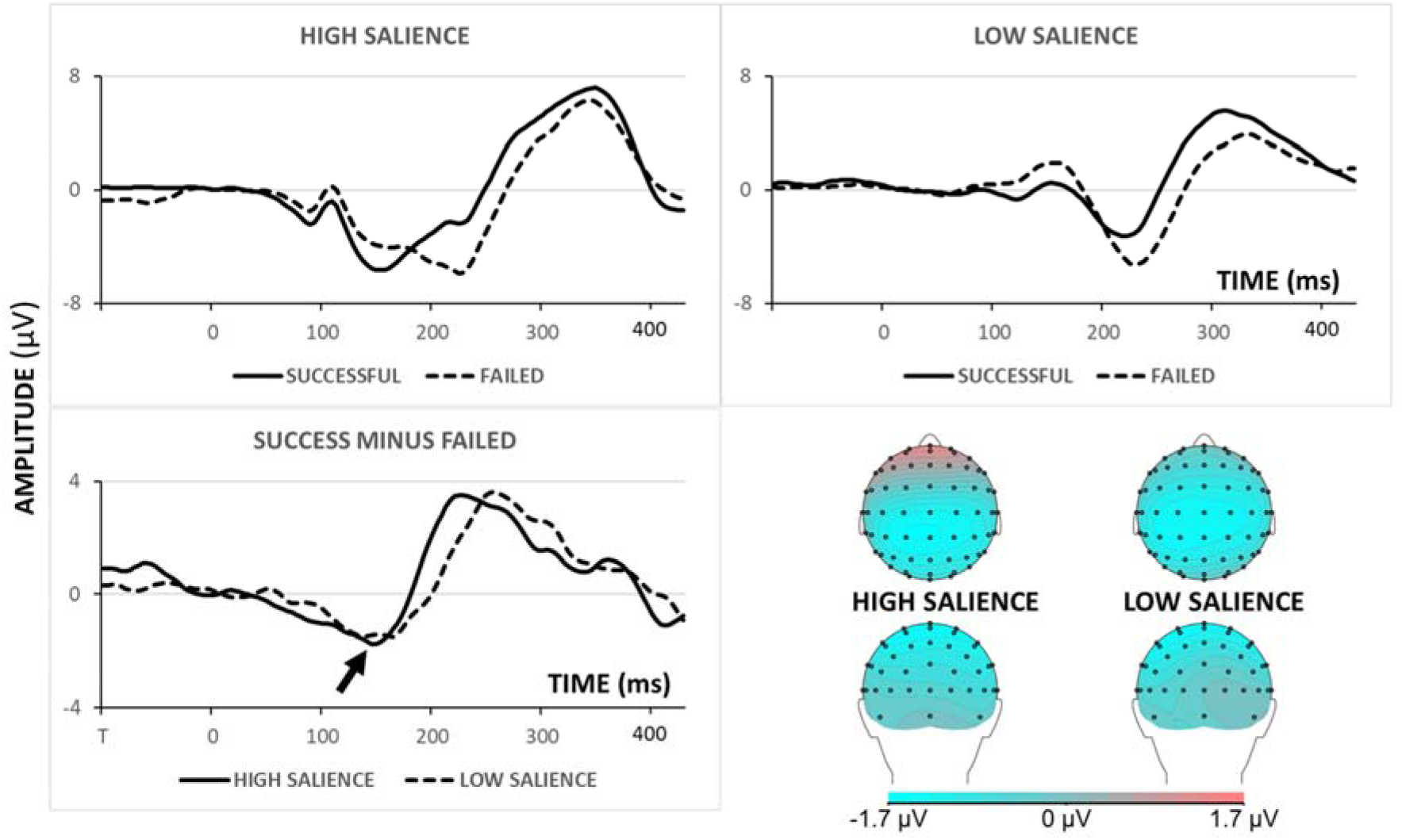
At electrode CPz: The stop-related negativity (arrow, Successful versus Failed stops), and scalp distribution (112-152 ms) for the visual high- and low-salience condition, respectively.

In Figure 4 it can also be seen that the stop-related negativity appears to evolve some tens of milliseconds before 100-ms latency. To accommodate this potential contamination from go-related activity, the significant Stop-Success effects in the collapsed 112-152 window were evaluated again using baseline values at 68-80 ms (the C1-like peak latency), as well as at 88-108 ms (the latency of the subsequent positive peak, see Figure 3). In all cases significance was retained, although p values became gradually larger with baseline more closely in time to the 112-ms latency (all *p* < .02).

#### Visual stop N2

The “*Search for visual stop-N2”* procedure outlined in the Method section revealed Stop-Success main effects for the right-frontal ROI (F6, F8, FC6) in the 196-212-ms time interval (F(1,28) = 17.4, *p* < .0005). As expected, this right-frontal stop N2 was especially pronounced in the low-salience condition (left arrow in Figure 5), where it featured a typical right-frontal scalp distribution (right arrow in Figure 5). However, statistically, the interaction between Salience and Stop Success effects was not at all significant (F(1,28) = 2.9, *p* = .1; BF10 = 0.72, anecdotal evidence for the null hypothesis). To further elucidate the absence of evidence for the alternative hypothesis, in the face of the apparent difference in Figure 5, we note that stop N2 was highly significant for low salience (F(1,28) = 20.1, *p* < .0005) but not for high salience (F(1,28) = 2.2). This implies that across participants stop N2 with low salience was highly systematic, but that only a fraction of these participants showed a substantial reduction with high salience.

**Figure 5.**
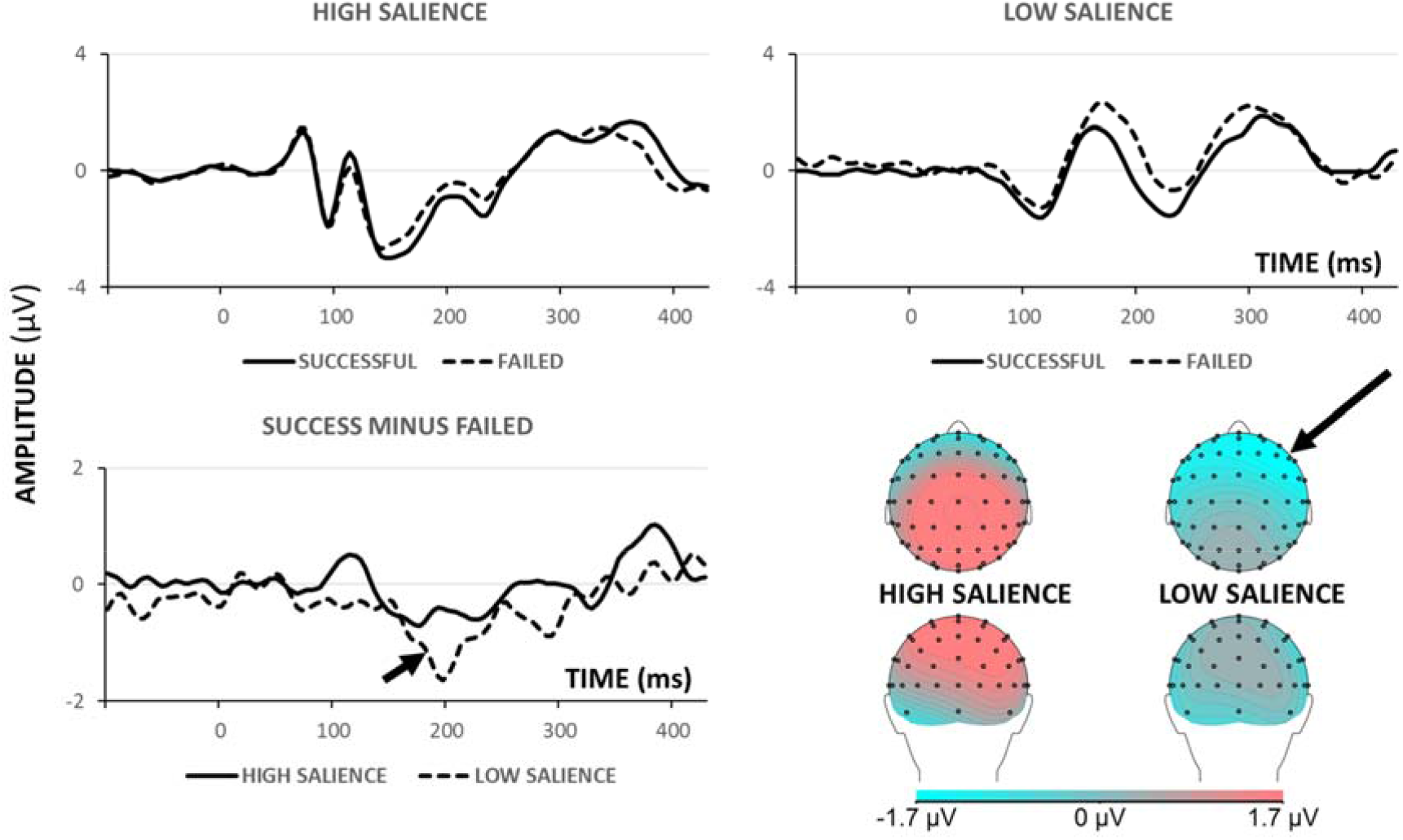
At the right-frontal ROI (average of F6, F8, FC6): The stop-related negativity (arrow, Successful versus Failed stops), and scalp distribution (196-212 ms) for the visual high- and low-salience condition, respectively.

#### Stop P3

Based on our previous work, on Wessel & Aron (2015), and on the current SSRT results, we expected stop-P3 (successful minus failed contrast) to peak before approximately 200 ms (auditory) and around 250 ms (visual-low salience), with peak latency for visual-high salience probably somewhere in between. Figure 7 confirms this expectation. Figure 8 depicts the corresponding waveforms.

**Figure 7.**
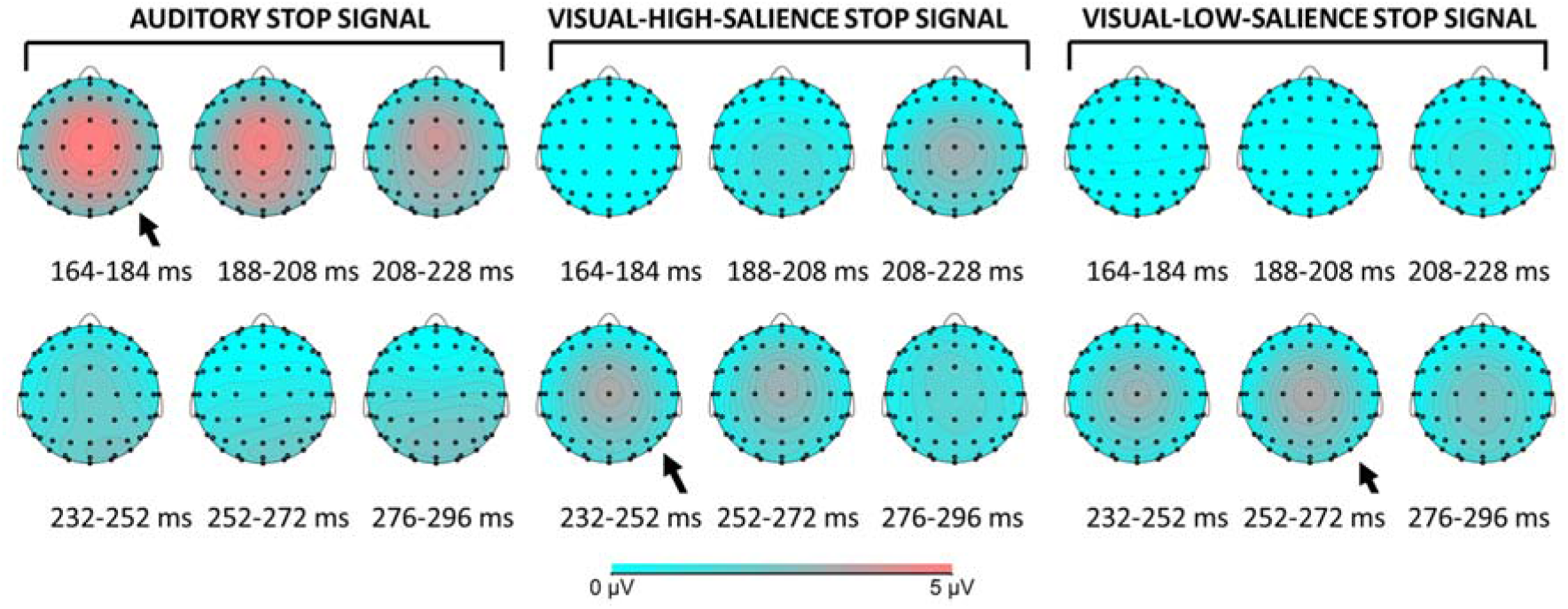
Scalp distributions of the Stop P3 (Successful minus Failed contrast) in the three stop-signal conditions, between 164 and 296 ms latency. Arrows indicate the apparent time windows of maximum amplitudes.

**Figure 8.**
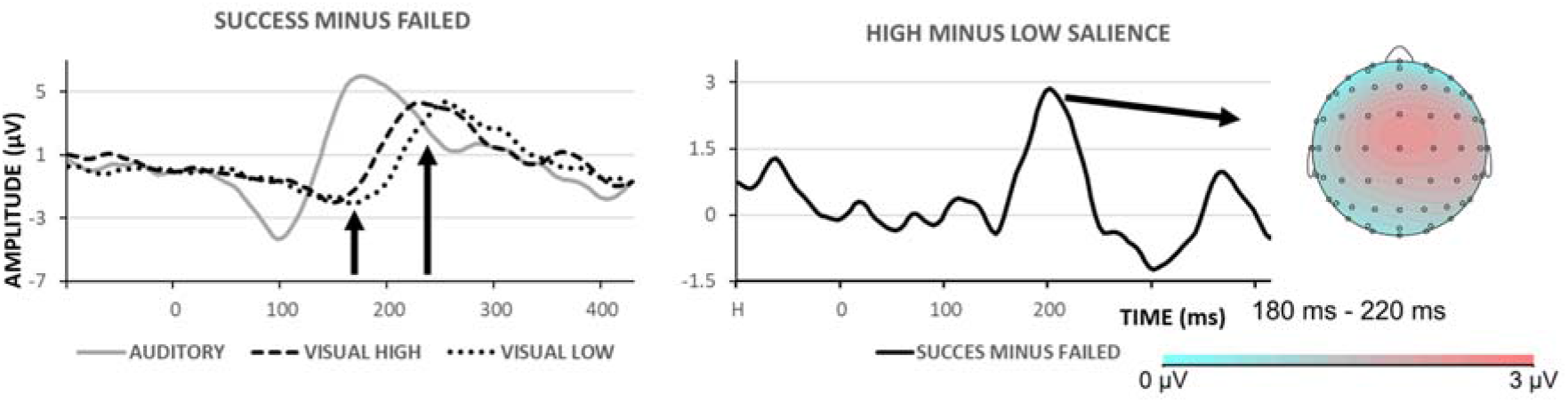
LEFT: Stop-P3 waveforms for the three stop-signal conditions. Arrows indicate the 180-220 ms time window for which the apparent latency difference for high versus low salience was tested. RIGHT: Difference (of difference waveforms) waveform highlighting the latency difference between high and low salience, and the corresponding scalp distribution.

For the auditory condition average amplitudes in the 160-208-ms latency window were significantly more positive for success than for failed: F(1,28) = 41.4, *p* < 0.001. Recently significant correlations between P3 amplitude separately for successful and failed stops and SSRT have been reported (Skippen et al., 2020). These results were not replicated for the present data, the correlation values were lower and even of reversed sign for failed stops (Pearson’s r = .26, reflecting larger P3 amplitudes not significantly associated with longer SSRTs; r = -.15 for successful stops). However, for the stop P3 proper (from successful-failed contrast) the correlation with SSRT was significant in the expected direction (r = -.591, *p* = .001; see Figure 9-left panel).

**Figure 9.**
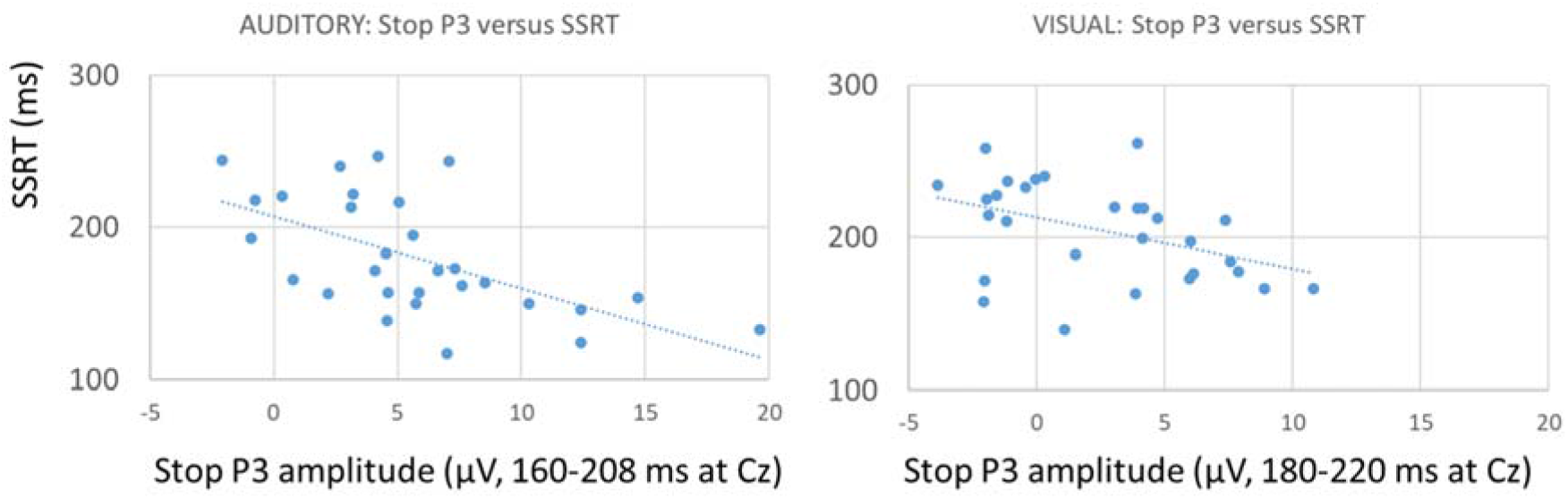
Auditory and visual-high-salience conditions: correlation between stop P3 and SSRT

Peak latencies for the visual conditions were longer than for auditory (omnibus Modality/ Salience effect: F(2,26) = 19.4, *p* <. 0001; see Figure 7), ranging from 206.2 ms (auditory), 251.4 ms (visual-high salience), and 264 ms (visual-low salience). Auditory differed from visual-low and from visual-high (both *p* < .001), but the difference between the two visual conditions was not confirmed (*p* = .25).

To zoom in better on the difference between the visual conditions especially after the onset of the stop P3 in the high-salient condition (see also Wessel & Aron, 2015), the differential onset was quantified as the average amplitude in the 180-220 ms time window. This apparent difference was confirmed statistically, F(1,28) = 11.1, *p* < .005. Specifically, this part of the stop P3 differed significantly from zero for high salience (F(1,28) = 11.4, *p* < .005), but not for low salience (F(1,28) = 1.1, *p* = .30). Also, the extent of this activity correlated significantly with SSRT for high salience (r= -.421, *p* < .025; see Figure 9-right panel), but not for the low-salience condition.

## Discussion

The present study addressed the effect of the salience of cues demanding suppression (stopping) of an ongoing response on behavioral and electro-cortical (ERP) measures of inhibitory control. As to the behavioral measures, stop-signal reaction time (SSRT) was shorter for auditory than for visual cues (stop signals), and shorter for high-salient than low-salient visual stop signals. Given this, which ERP measure, defined as differential activity for successful versus failed stops responds in a similar way to the salience manipulation: stop N1, stop N2, and/ or stop P3? Robust effects of salience were observed only for stop P3, of which the onset latencies varied across modality and visual salience in a manner similar to that for SSRT. Specifically, in the 180-220-ms latency interval, stop P3 was significant for high but not for low salience. Explicit support for a functional relation between this stop P3 and SSRT was their significant correlation in the high-salience condition: The more pronounced the stop P3 in this interval, the shorter the SSRT.

An explicit aim of the present study was to identify a potential visual stop N1 (100-200 ms latency) and the visual stop N2 (right frontal, around 200 ms latency) in one sample of healthy subjects. Indeed, in the present visual stop-signal conditions a relatively early (112-152 ms) deflection was observed that was stronger for successful than for failed stop (visual “stop N1”), and additionally a later (196-212 ms) negativity which followed the same successful/ failed pattern and was at maximum above right-frontal area’s (visual “stop N2”). However, our prediction that the visual stop N1 would be stronger for high-salient visual stop signals, and the visual stop N2 for low-salient ones, was not at all confirmed. It should be noted that Bayes factors indicated moderate and perhaps positive evidence for the null hypotheses (regarding the salience x stop success interaction) for visual stop N1, but that for the stop N2 the evidence was inconclusive, with the interaction going in the expected direction. As explained in the section on participants, for visual stop N1 BF01 increased with an increasing number of subjects. However, a posteriori analysis for stop N2 revealed a BF01 of 3.88 with N=20 and BF01 = 1.39 with N=29, indicating that a larger sample size might allow to detect a sensitivity for visual salience for stop N2 after all.

It has been speculated that a visual stop N1, analogous to the auditory stop N1, reflects activity in visual cortex (100-200 ms latency) especially related to the detection of rare events, such as an infrequent stop signal (Kenemans et al., 2003; Kenemans, 2015; Kimura et al., 2009). This idea leads to the expectation of visual-stop-N1 scalp distribution across occipital area’s. The currently observed CPz maximum does, at least qualitatively, not support this idea. Admittedly, this topography does not exclude an intracranial generator in occipital areas; but the pattern is quite different from the sharp focus over occipital area’s that has been consistently reported for the rareness-related negativity (RRN, e.g., Kenemans et al., 2003, 2010; Kimura et al., 2009). The currently observed centro-parietal midline focus could as well be consistent with what has been labeled a non-lateralized stop-related fronto-central N2 that precedes the stop P3 in time (Huster et al., 2020; Wessel & Aron, 2015). Of course a more definitive answer to this question should result from comparing RRN and visual stop N1 in the same sample of subjects.

The currently observed right-frontal distribution for the stop N2 was as predicted, and consistent with the earlier findings reported by Schmajuk et al. (2006). As noted in the introduction, it is possible that this topography reflects activity in right inferior frontal gyrus induced by visual stop signals that is stronger in case of successful stops. Again, at least as far as ERPs reveal, such right-frontal activity was not seen in response to auditory stop signals. These were marked by the auditory stop N1 immediately followed by the stop P3. Both stop N1 and stop P3 have been frequently reported before; for stop N1 this work is about the eightest study in which it was robustly observed (see references in Introduction), and for stop P3 this number is undoubtedly much higher.

The current analysis also addressed the pre-100ms-latency visual evoked potential (VEP) that could be expected especially for the high-salient visual condition. Indeed, a pronounced negative peak evolved and dissolved between 50 and 100 ms with a clear topographical focus over medial occipital cortex, specifically for the high-salient condition. This provides evidence that the current salience manipulation indeed resulted in stronger activity in visual cortex. It has been convincingly argued that such short-latency medial-occipital negativities originate from primary visual cortex (e.g., Kenemans et al., 2000). Thus, the presently observed occipital focus can be viewed as both an anchor for and a validation of the topographical information with respect to the stop-related processes. Note that anecdotical (inconclusive, BF10 = 0.53) evidence was found against the hypothesis that this VEP was associated with stopping success. For the visual stop signals, it is rather after 100 ms latency that the first significant association with stopping success was observed (visual stop N1).

It may be noted that salience is also influenced by the extent of perceptual overlap between stop signal and go context. Multiple-resource theory postulates limited capacities in human information tied to the amount of perceptual overlap between concurrent streams of information (Wickens, 2008). For visual go signals, an auditory stop signal implies a virtual absence of perceptual overlap. A tonically potentiated inhibitory connection between the auditory and motor cortex would therefore not interfere with activating signals from letter representations in visual cortex.

A potentially interesting by-catch in the present data was the partial replication of auditory N1 and P3 amplitude correlations with SSRT separately for successful and failed stops as reported recently by Skippen et al. (2020): Larger negative N1s are associated with shorter SSRTs. In contrast the stop-N1 proper did not show such a correlation (this correlation was not discussed in Skippen et al.). For the auditory P3 the replication of the Skippen et al. results failed (for successful and failed stops separately); but here the correlation for the stop P3 was in fact observed (larger stop P3s associated with shorter SSRTs), and this also held for the stop P3 in the high-salient visual condition. It is beyond the scope of this article, and perhaps also premature, to present a full mechanistic account for these differential correlation patterns. For the time being we present two further remarks. First, the differential correlation patterns seem to be consistent with results reported by Lansbergen et al. (2007), who compared healthy slow and fast stoppers (long versus short SSRTs), and found smaller stop P3s for the fast stoppers, but no difference in stop N1 (separate analyses for successful and failed were not reported). It is also worth noting that similar differences between fast and slow stoppers or stopping have been reported for fMRI-assessed activation in the presupplementary motor area, the putative source of the stop P3 (Chao et al., 2006; Duann et al., 2009; Li et al., 2009; see Kenemans, 2015, for review). Second, the differential correlation pattern for stop N1 versus stop P3 embodies a further double dissociation between the two processes, that adds to the dissociation in terms of individual differences (Lansbergen et al., 2007), and in terms of pharmacological effects (stop N1 in ADHD is restored by methylphenidate, but stop P3 is not; Overtoom et al., 2009).

A final remark should be made in relation to recent computational modelling studies that address the role of potential “trigger failures” (TFs): On a minority of stop trials, the stopping process is simply not activated (see Skippen et al., 2020, for an overview). Briefly, these studies indicate that including TFs in the model results in fairly shorter estimates for SSRT (e.g., for auditory SSRT 196.9 ms without TFs versus 132.4 with TFs included, Skippen et al., 2020). Furthermore, also TF rate is correlated with N1 and P3 amplitudes in an intuitive manner (larger amplitudes associated with lower TF rates). Therefore it might be that the presently reported correlations between SSRT and N1 and P3 actually reflect a mixture of correlations with TF-corrected SSRT and TF rate. An about 60 ms shorter estimate for SSRT would also pose complications for a potential causal relation between stop P3 and SSRT, given the then clearly reversed temporal order. In that scenario it is possible that stop P3 reflects suppression of later phases of peripheral and central motor activation (cf. De Jong et al., 1990).

To conclude, the present study revealed that an across- and within-modality manipulation of salience resulted in gradual variation of inferred stopping speed. The only likely electro-cortical mechanism revealed in the present analysis to parallel this gradual variation was the onset of the stop P3. Shorter-latency stop-related mechanisms were also observed but did not vary as a function of salience. To the extent that shorter-latency mechanisms reflect more proactive, and stop P3 more reactive inhibitory control, it is the reactive mechanism that may be involved in salience-induced enhancement of inhibitory control.

## Author contributions

J.L. Kenemans: Conceptualization; Methodology; Formal analysis; Writing

I. Schutte: Data acquisition; Formal analysis; Review

S. van Bijnen: Data acquisition; Formal analysis; Review

H.N.A. Logemann: Conceptualization; Methodology; Formal analysis; Writing; Review

## Acknowledgements

Remo van der Heiden, assistance in formal analysis

Koen Böcker and Merlijn Hutteman, initial discussion on conceptualization

## Notes

### Competing Interest Statement

The authors have declared no competing interest.

## References

Aron, A. R., Fletcher, P. C., Bullmore, E. T., Sahakian, B. J., & Robbins, T. W. (2003). Stop-signal inhibition disrupted by damage to right inferior frontal gyrus in humans. Nat Neurosci, 6(2), 115–116.

Bekker, E. M., Kenemans, J. L., Hoeksma, M. R., Talsma, D., & Verbaten, M. N. (2005). The pure electrophysiology of stopping. Int J Psychophysiol, 55(2), 191–198. doi: S0167-8760(04)00163-1 [pii]10.1016/j.ijpsycho.2004.07.005

Bekker, E. M., Overtoom, C. C., Kooij, J. J., Buitelaar, J. K., Verbaten, M. N., & Kenemans, J. L. (2005). Disentangling deficits in adults with attention-deficit/hyperactivity disorder. Arch Gen Psychiatry, 62(10), 1129-1136. doi: 62/10/1129 [pii]10.1001/archpsyc.62.10.1129

Bekker, E. M., Overtoom, C. C. E., Kooij, J. J. S., Buitelaar, J. K., Verbaten, M. N., & Kenemans, J. L. (2005). Disentangling Deficits in Adults With Attention-Deficit/Hyperactivity Disorder. Arch Gen Psychiatry, 62(10), 1129–1136.

Blizzard, S., Fierro-Rojas, A., & Fallah, M. (2016). Response Inhibition Is Facilitated by a Change to Red Over Green in the Stop Signal Paradigm. Front Hum Neurosci, 10, 655. doi:10.3389/fnhum.2016.00655

Boehler, C. N., Munte, T. F., Krebs, R. M., Heinze, H. J., Schoenfeld, M. A., & Hopf, J. M. (2009). Sensory MEG responses predict successful and failed inhibition in a stop-signal task. Cereb Cortex, 19(1), 134–145. doi: bhn063 [pii]10.1093/cercor/bhn063

Chao, H., Luo, X., Chang, J., & Li, C.-s. (2009). Activation of the pre-supplementary motor area but not inferior prefrontal cortex in association with short stop signal reaction time - an intra-subject analysis. BMC Neuroscience, 10(1), 75.

De Jong, R., Coles, M. G., & Logan, G. D. (1995). Strategies and mechanisms in nonselective and selective inhibitory motor control. J Exp Psychol Hum Percept Perform, 21(3), 498–511.

De Jong, R., Coles, M. G., Logan, G. D., & Gratton, G. (1990). In search of the point of no return: the control of response processes. J Exp Psychol Hum Percept Perform, 16(1), 164–182.

Duann, J.-R., Ide, J. S., Luo, X., & Li, C.-s. R. (2009). Functional Connectivity Delineates Distinct Roles of the Inferior Frontal Cortex and Presupplementary Motor Area in Stop Signal Inhibition. The Journal of Neuroscience, 29(32), 10171-10179. doi:10.1523/jneurosci.1300-09.2009

Goss-Sampson, M., Van Doorn, J., & Wagenmakers, E. J. (2020). Bayesian Inference in JASP: A New Guide For Students: http://static.jasp-stats.org/Manuals/Bayesian_Guide_v0_12_2_1.pdf.

Gratton, G., Coles, M. G., & Donchin, E. (1983). A new method for off-line removal of ocular artifact. Electroencephalogr Clin Neurophysiol, 55(4), 468–484.

Hughes, M. E., Fulham, W. R., Johnston, P. J., & Michie, P. T. (2012). Stop-signal response inhibition in schizophrenia: Behavioural, event-related potential and functional neuroimaging data. Biological Psychology, 89(1), 220-231. doi:https://doi.org/10.1016/j.biopsycho.2011.10.013

Huster, R. J., Messel, M. S., Thunberg, C., & Raud, L. (2020). The P300 as marker of inhibitory control – Fact or fiction? Cortex, 132, 334-348. doi:https://doi.org/10.1016/j.cortex.2020.05.021

Kenemans, J. L. (2015). Specific proactive and generic reactive inhibition. Neuroscience & Biobehavioral Reviews, 56(0), 115-126. doi: http://dx.doi.org/10.1016/j.neubiorev.2015.06.011

Kenemans, J. L., Baas, J. M. P., Mangun, G. R., Lijffijt, M., & Verbaten, M. N. (2000). On the processing of spatial frequencies as revealed by evoked-potential source modeling. Clinical Neurophysiology, 111(6), 1113–1123.

Kenemans, J. L., Grent-’t Jong, T., & Verbaten, M. N. (2003). Detection of visual change: Mismatch or rareness? NeuroReport., 14, 1239–1243.

Kenemans, J. L., Hebly, W., van den Heuvel, E., & Grent-’T-Jong, T. (2010). Moderate alcohol disrupts a mechanism for detection of rare events in human visual cortex. Journal of Psychopharmacology, 24(6), 839–845. doi:10.1177/0269881108098868

Kimura, M., Katayama, J., Ohira, H., & Schröger, E. (2009). Visual mismatch negativity: New evidence from the equiprobable paradigm. Psychophysiology, 46(2), 402–409.

Lansbergen, M. M., Bocker, K. B., Bekker, E. M., & Kenemans, J. L. (2007). Neural correlates of stopping and self-reported impulsivity. Clin Neurophysiol, 118(9), 2089–2103. doi: S1388-2457(07)00302-1 [pii]10.1016/j.clinph.2007.06.011

Li, C.-s., Huang, C., Constable, R. T., & Sinha, R. (2006). Imaging Response Inhibition in a Stop-Signal Task: Neural Correlates Independent of Signal Monitoring and Post-Response Processing. The Journal of Neuroscience, 26(1), 186–192. doi:10.1523/jneurosci.3741-05.2006

Matzke, D., Hughes, M., Badcock, J. C., Michie, P., & Heathcote, A. (2017). Failures of cognitive control or attention? The case of stop-signal deficits in schizophrenia. Attention, Perception, & Psychophysics, 79(4), 1078–1086. doi:10.3758/s13414-017-1287-8

Montanari, R., Giamundo, M., Brunamonti, E., Ferraina, S., & Pani, P. (2017). Visual salience of the stop-signal affects movement suppression process. Exp Brain Res, 235(7), 2203–2214. doi:10.1007/s00221-017-4961-0

Näätänen, R., & Picton, T. (1987). The N1 wave of the human electric and magnetic response to sound: a review and an analysis of the component structure. Psychophysiology, 24(4), 375–425.

Overtoom, C. C., Bekker, E. M., van der Molen, M. W., Verbaten, M. N., Kooij, J. J., Buitelaar, J. K., & Kenemans, J. L. (2009). Methylphenidate restores link between stop-signal sensory impact and successful stopping in adults with attention-deficit/hyperactivity disorder. Biol Psychiatry, 65(7), 614–619. doi: S0006-3223(08)01434-0 [pii]10.1016/j.biopsych.2008.10.048

Pliszka, S. R., Liotti, M., Bailey, B. Y., Perez, R., 3rd, Glahn, D., & Semrud-Clikeman, M. (2007). Electrophysiological effects of stimulant treatment on inhibitory control in children with attention-deficit/hyperactivity disorder. J Child Adolesc Psychopharmacol, 17(3), 356–366. doi: 10.1089/cap.2006.0081

Pliszka, S. R., Liotti, M., & Woldorff, M. G. (2000). Inhibitory control in children with attention-deficit/hyperactivity disorder: event-related potentials identify the processing component and timing of an impaired right-frontal response-inhibition mechanism. Biol Psychiatry, 48(3), 238–246. doi: S0006-3223(00)00890-8 [pii]

Polich, J. (2007). Updating P300: An integrative theory of P3a and P3b. Clinical Neurophysiology, 118(10), 2128–2148. doi: http://dx.doi.org/10.1016/j.clinph.2007.04.019

Schmajuk, M., Liotti, M., Busse, L., & Woldorff, M. G. (2006). Electrophysiological activity underlying inhibitory control processes in normal adults. Neuropsychologia, 44(3), 384–395. doi: S0028-3932(05)00213-7 [pii]10.1016/j.neuropsychologia.2005.06.005

Skippen, P., Fulham, W. R., Michie, P. T., Matzke, D., Heathcote, A., & Karayanidis, F. (2020). Reconsidering electrophysiological markers of response inhibition in light of trigger failures in the stop-signal task. Psychophysiology, 57(10), e13619. doi:10.1111/psyp.13619

Tannock, R., Schachar, R. J., Carr, R. P., Chajczyk, D., & Logan, G. D. (1989). Effects of methylphenidate on inhibitory control in hyperactive children. J Abnorm Child Psychol, 17(5), 473–491.

van der Schoot, M., Licht, R., Horsley, T. M., & Sergeant, J. A. (2005). Effects of stop signal modality, stop signal intensity and tracking method on inhibitory performance as determined by use of the stop signal paradigm. Scand J Psychol, 46(4), 331–341. doi:10.1111/j.1467-9450.2005.00463.x

Verbruggen, F., Aron, A. R., Band, G. P., Beste, C., Bissett, P. G., Brockett, A. T., … Boehler, C. N. (2019). A consensus guide to capturing the ability to inhibit actions and impulsive behaviors in the stop-signal task. Elife, 8. doi:10.7554/eLife.46323

Wessel, J. R., & Aron, A. R. (2013). Unexpected Events Induce Motor Slowing via a Brain Mechanism for Action-Stopping with Global Suppressive Effects. The Journal of Neuroscience, 33(47), 18481–18491. doi: 10.1523/jneurosci.3456-13.2013

Wessel, J. R., & Aron, A. R. (2014). It’s not too late: The onset of the frontocentral P3 indexes successful response inhibition in the stop-signal paradigm. Psychophysiology, n/a-n/a. doi: 10.1111/psyp.12374

Wessel, J. R., & Aron, A. R. (2017). On the Globality of Motor Suppression: Unexpected Events and Their Influence on Behavior and Cognition. Neuron, 93(2), 259–280. doi: https://doi.org/10.1016/j.neuron.2016.12.013

Wickens, C. D. (2008). Multiple resources and mental workload. Hum Factors, 50(3), 449–455.

Woldorff, M. G. (1993). Distortion of ERP averages due to overlap from temporally adjacent ERPs: analysis and correction. Psychophysiology, 30(1), 98–119.

